# deGSM: memory scalable construction of large scale de Bruijn Graph

**DOI:** 10.1101/388454

**Authors:** Hongzhe Guo, Yilei Fu, Yan Gao, Junyi Li, Yadong Wang, Bo Liu

## Abstract

**Motivation:** De Bruijn graph, a fundamental data structure to represent and organize genome sequence, plays important roles in various kinds of sequence analysis tasks such as de novo assembly, high-throughput sequencing (HTS) read alignment, pan-genome analysis, metagenomics analysis, HTS read correction, etc. With the rapid development of HTS data and ever-increasing number of assembled genomes, there is a high demand to construct de Bruijn graph for sequences up to Tera-base-pair level. It is non-trivial since the size of the graph to be constructed could be very large and each graph consists of hundreds of billions of vertices and edges. Current existing approaches may have unaffordable memory footprints to handle such a large de Bruijn graph. Moreover, it also requires the construction approach to handle very large dataset efficiently, even if in a relatively small RAM space.

**Results:** We propose a lightweight parallel de Bruijn graph construction approach, de Bruijn Graph Constructor in Scalable Memory (deGSM). The main idea of deGSM is to efficiently construct the Bur-rows-Wheeler Transformation (BWT) of the unipaths of de Bruijn graph in constant RAM space and transform the BWT into the original unitigs. It is mainly implemented by a fast parallel external sorting of k-mers, which allows only a part of k-mers kept in RAM by a novel organization of the k-mers. The experimental results demonstrate that, just with a commonly used machine, deGSM is able to handle very large genome sequence(s), e.g., the contigs (305 Gbp) and scaffolds (1.1 Tbp) recorded in GenBank database and Picea abies HTS dataset (9.7 Tbp). Moreover, deGSM also has faster or comparable construction speed compared with state-of-the-art approaches. With its high scalability and efficiency, deGSM has enormous potentials in many large scale genomics studies.

**Availability:** https://github.com/hitbc/deGSM.

**Contact:** (YW) and (BL)

**Supplementary information:** Supplementary data are available online.

## 1 Introduction

De Bruijn graph is a fundamental data structure to represent and organize genome sequences. It is widely used in many sequence analysis tasks such as de novo genome assembly (Zerbino et al., 2008; Gnerre et al., 2011; Luo et al. 2012), high-throughput sequencing (HTS) read alignment (Liu et al., 2016; Siren, 2017), pan-genome analysis (Marcus et al., 2014), metagenomics species classification (Guan et al., 2016), transcript isoform identification and quantification (Bray et al., 2016) and HTS read correction (Salmela et al., 2017; Limasset, A. et al., 2018), etc.

HTS is being ubiquitously applied in the field of genomics, supporting many large scale genome research projects on human populations and various species, and generating massive HTS datasets continuously. Currently, sequencing a large genome can produce a dataset in up to Tera base pairs (bps), e.g., the whole genome sequencing (WGS) dataset of the 20 Gbp Picea abies genome is as large as 9.7 Tera bps. Moreover, the size of the datasets for population sequencing can be even larger. Meanwhile, as the number of assembled genomes continues to rise, there have already been more than one assembled genomes for each specific species. These genomes are directly used for pan-genome analysis, or used as reference sequences for various tasks like read alignment (Liu et al., 2016), variant calling (Eggertsson et al., 2017), species classification (Guan et al., 2016), etc. However, the total size of these sequences is enormous, e.g., the sizes of contigs and scaffolds recorded in Genbank are over 305 Giga bps and

### 1.1 Tera bps, respectively

De Bruijn graph has various applications on the analysis of these data. However, the construction of de Bruijn graph could be a major bottleneck due to two issues: 1) the large input sequence itself could lead to a huge number of the vertices; 2) the potential sequencing error of HTS datasets can make the number of vertices of the graph explosively increase. Under such circumstance, the size of the graph to be constructed would be very large and the graph construction approach needs to be highly scalable to adapt to commonly used computational environments.

Many efforts have been made in the construction of de Bruijn graph. Basically, the proposed approaches can be categorized as hash-based and suffix-trie-based.

Hash-based approaches generally build hash tables to represent and organize the vertices of de Bruijn graph. This approach is frequently used in de novo assembly (Zerbino et al., 2008; Gnerre et al., 2011; Luo et al. 2012). In this category, two hash-based techniques, Minimizer and Bloom filter are widely used.

Minimizer is the representative sequence selected from a group of adjacent k-mers which have identical substring (Roberts et al., 2004; Wood and Salzberg, 2014). It is often used as seed to reduce the storage of k-mers in hash table. Movahedi et al (Movahedi. et al., 2012) used minimizer during de Bruijn graph construction to partition de Bruijn graph into slices. This helps to reduce the memory footprint of de novo assembly. Chikhi et al (Chikhi et al., 2014) proposed a novel algorithm, BCALM, which is based on frequency-based minimizer instead of the commonly used lexicographic order-based minimizer. Meanwhile, this work proposed the DBGFM data structure which also takes advantage of FM-index to represent and compact the output of BCALM. Further, Chikhi et al (Chikhi et al., 2016) took advantage of a novel partitioning strategy of frequency-based minimizer to achieve better parallelization. It demonstrates that the proposed approach has the ability to construct the graph for the HTS datasets of large genomes, e.g., >20 Gbp pine and spruce genomes.

Bloom filter is a space-efficient probabilistic data structure that is used to check the presence of an element. This data structure can largely reduce the size of the graph due to its bit array-based design. However, false positives also could be introduced into the data structure due to probabilistic design. So specific methods are necessary to handle the false positives, usually with an extra cost of RAM space. Pell (Pell et al., 2012) used Bloom filter to store vertices and partition the whole graph into components, which reduces the memory footprint of metagenome assembly. The experimental results indicate that there are about 15% false positive bits in Bloom filter-based data structure. Chikhi and Rizk (Chikhi and Rizk, 2013) proposed a novel marking scheme along with an auxiliary structure to enumerate specific vertices, and Drezen et al(Drezen et al., 2014) demonstrated that this method is helpful in de novo assembly. Holley (Holley et al., 2015) proposed Bloom filter tree to index and query pangenome datasets. El-Metwally et al (El-Metwally et al., 2016) proposed a light-weight genome assembler using pattern-blocked Bloom filter to compress *k*-mers. TwoPaCo (Minkin et al., 2017) represents the non-branching paths (unipaths) of the graph with single edges, and uses bloom filter to compactly store them. It achieves a small memory footprint during graph construction, which supports to build de Bruijn graph for large genomes on a modern server.

Other Bit array-based data structures are also used in de Bruijn graph construction. For example, Conway and Bromage (Conway and Bromage, 2011) represents and constructs de Bruijn graph by a compressed bit vector initially proposed by (Okanohara and Sadakane, 2007).

Suffix trie-based data structures, such as suffix tree, Burrows-Wheeler Transformation (BWT), Ferragina-Manzini index (FM-index) (Ferragina and Manzini, 2000) and suffix array (Manber and Myers, 1993), are also used in de Bruijn graph construction. (Bowe et al., 2012) is the first effort to take advantage of XBW-transform to represent de Bruijn graph. The proposed data structure, succinct de Bruijn graph, has outstanding ability to compact de Bruijn graph, due to that the strings implied by the unipaths/unitigs can be succinctly represented. (R-land,2013) proposed a similar approach, named as KFM-index, which represents de Bruijn graph by the FM-index of the k-mers of the input sequence. Additionally, it is theoretically approved that the RAM usage can be reduced by keeping only a part of the graph in memory during construction. However, it does not provide an implementation which can work on large datasets. Li et al (Li et al. 2015) takes advantage of succinct de Bruijn graph to develop a metagenome assembler named as MEGAHIT. In this approach, both GPU-based and CPU-accelerated in-memory sorting approaches are used to construct the succinct de Bruijn graph of metagenome HTS dataset. This implementation has fast speed, but its memory footprint is large due to the in-memory design. Moreover, Marcus et al (Marcus et al., 2014) proposed a suffix tree-based approach, SplitMEM, to organize de Bruijn graph for pan-genome analysis. In this approach, a novel data structure, suffix skip, is introduced to facilitate the traversal of suffix links. It helps to efficiently decompose maximal exact matches into graph vertices, which is beneficial to pan-genome analysis. By taking advantage of the topological relationships between suffix trees and compressed de Bruijn graphs, SplitMEM can construct de Bruijn graph in linear time and space. Further, Baier et al (Baier et al., 2016) improves the time and space cost of SplitMEM with BWT and compressed suffix tree.

The construction of very large de Bruijn graph could be widely used in many ongoing and forthcoming large scale genomic studies. However, there is still lack of tools with good ability to construct de Bruijn graph consisting of tens to hundreds of billions of vertices in commonly used computational environment with fast speed. Such approach is necessary to properly handle large datasets for both HTS data and assembled genomes. Herein, we propose de Bruijn Graph Constructor in Scalable Memory (deGSM), a highly scalable approach suitable for constructing very large de Bruijn graph. DeGSM is a suffix-trie-based approach, in theory similar to succinct de Bruijn graph. Instead of directly constructing the graph itself, deGSM constructs the BWT of the unitigs of the graph and recovers the unitigs with the constructed BWT string. Specifically, deGSM implements a fast parallel external sorting of the *k*-mers in the graph to build the BWT of unitigs. The implementation of the external sorting can fully consider the CPU and RAM configurations and improve the speed with limited computational resources. More importantly, a novel organization of the *k*-mers is used to always allow to keep only a part of *k*-mers in RAM, which is critical to achieve constant RAM space cost. DeGSM can be scalable to construct de Bruijn graph for the HTS dataset of a large genome (e.g., the 20 Gbp Picea abies genome), or all the contigs or scaffolds (upto 1.1 Tbp) recorded in GenBank with 16GB or less RAM. Meanwhile, deGSM also has faster or comparable construction speed compared with state-of-the-art approaches.

## 2 Methods

### 2.1 Preliminary

Let a genome, S, be a sequence over the alphabet Σ = {*A, C, G, T*}. The de Bruijn graph of *S*, D, is a directed graph, where the vertices consist of all the *k*-mers of S. For any pair of vertices of D, (*V*_i_, *V*_j_), there is a directed edge *V*_i_ → *V*_j_, if the *k*-1 suffix of *V*_i_ is same to the *k*-1 prefix of *V*_j_. A set of maximal non-branched paths (unipaths) can be derived from D. Each of the unipaths meets the following conditions: i) for the first vertex, the in-degree is 0 or >1, and the out-degree is 1; ii) for the last vertex, the out-degree is 0 or >1, and the in-degree 1; iii) for all the other vertices along the path, the in- and out-degrees are exactly 1. This definition follows previous studies (Tomescu and Medvedev, 2016), and the string implied by compacting a certain unipath is called a “unitig” (Gnerre *et al.*, 2011; Zimin et al., 2013).

For a given string T defined on Σ, the BWT of T$ is defined as the permutation of the characters of T that, *B*_T_[*i*] = *T*[*SA*[*i*] - 1], if *SA*[*i*] ≠ 0, and *B_T_*[*i*] = Σ, otherwise, where *SA*[*i*] indicates the starting position of the *i*-th lexicographically smallest suffix of T, and Σ is an auxiliary character lexicographically larger than any character of Σ.

Let the set of the unitigs of D be *U* = {*U_i_*, *i* = 1, …, |*U*|}, where |*U*| is the number of the unitigs, and a string C = *U*_1_#*U*_2_# … #*U*_|U|_Σ is a specific permutation of the unitigs, where # is an auxiliary character lexicographically larger than any character of Σ, but smaller than Σ. D can be compactly represented by the BWT of C (indicated as *B_C_*). Herein, we term this representation as “unitig-BWT”, which is to some extent similar to the data structure used in DBGFM (Chikhi et al., 2014).

*B_C_* can be used as a self-index of D, i.e., all the vertices of D can be accessed by querying *B_C_* for the corresponding *k*-mers, and all the in- and out-edges of a given vertex can be accessed by querying *B_C_* for the *k*-mers corresponding to the connected vertices. It is also worthnoting that the edge query operation is only needed for the end vertices of unipaths when traversing D, since the traversing on the internal vertices of unipaths can be efficiently done by the LF-mapping operation of BWT (Ferragina and Manzini, 2000). Moreover, all the original unitigs can be directly derived from *B_C_* in linear time with the LF-mapping.

Two additional definitions on C are given as following:

*k^>^-prefix*: for any suffix of C starting from a position ≥ *k* characters away from ‘#’, the corresponding *k^>^-prefix* is the length *k* prefix of the suffix. It is obvious that the *k^>^-prefixes* are the vertices (*k*-mers) of D.

*k^<^-prefix*: for any suffix of C starting from a position < *k* characters away from ‘#’, the corresponding *k^<^-prefix* is the prefix of the suffix ending at the next *k*-mer, if the length of the suffix is longer than *k*+1, or suffix itself, otherwise.

A schematic illustration of these concepts is in Fig. 1.

**Fig. 1:**
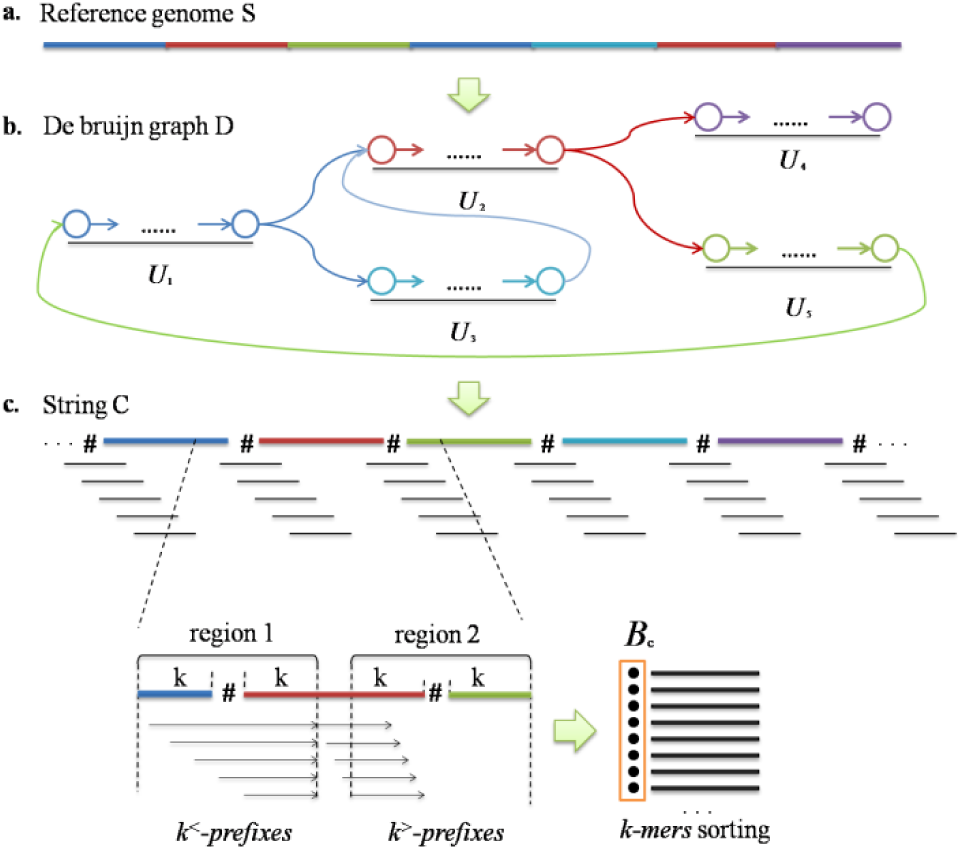
A schematic illustration of the deGSM method. (a) There is a large number of repetitive fragments in the reference genome sequence or sequencing data (strips with various colors). Here the repetition represented by blue strip has two copies et.al. (b) The copies of each repetitive sequence collapse into every uni-path of de Bruijn graph. (c) Make a specific permutation of all unipaths to form string *C* and different unipaths are separated by separator #. The *k^<^-prefix* can be generated from a local region every other 1 bp with the maximum length of 2*k*. The *k^>^-prefix* with a length of *k* starts at the position ≥*k* characters away from ‘#’. Under such circumstance, each *k^<^-prefix* has character # while the *k^>^-prefix* does not cross any separator. Merge *k^<^-prefixes* and *k^>^-prefixes* to generate *B_C_*

There are three characteristics of *k^>^-prefix* and *k^<^-prefix* as following, which are critical to the construction of *B_C_*.

1) Since all the vertices (*k*-mers) of D are distinct, the lexicographical orders of any two suffixes of C can be determined by only considering the corresponding *k^>^-prefix* or *k^<^-prefix*. Thus,*B_C_* can be constructed by sorting *k^>^-prefixes* and *k^<^-prefixes*.

2) C can be presented by the various permutations of the unitigs, but the *k^>^-prefix* set is irrelevant to the permutation, since *k^>^-prefixes* are exactly the vertices of D. Under such circumstance, the *k^>^-prefix* set can be obtained by only constructing the vertex set of D.

3) Any *k^<^-prefix* can be seen as a concatenation between the end vertex of one unipath and the starting vertex of the next unipath, and this depends on the permutation of the unitigs. However, it is worthnoting that *k^<^-prefixes* only depends on the permutation of the vertices at the two ends of the unitigs (Fig.1), which can be straightforwardly made with only the vertices at the two ends of the unipaths. In this situation, the *k^<^-prefix* set can be obtained by only identifying those end vertices, regardless of constructing the whole unitigs.

These characteristics make it possible to construct *B_c_* by the construction and sorting of *k^>^-prefixes* and *k^<^-prefixes*. This is helpful to handle large genome sequence since all the steps can be implemented with constant space cost. Specifically, deGSM takes advantage of these characteristics to make a lightweight and efficient de Bruijn graph construction.

### 2.2 Overview of deGSM approach

DeGSM constructs the *B_C_* representation of the de Bruijn graph D for a given genome S in five steps as following (a flowchart is in Fig.2):

1) recognizing all distinct (*k*+2)-mers of S;

2) sorting the (*k*+2)-mers in lexicographical order;

3) using the sorted (*k*+2)-mers to build a sorted table of the vertices of D (i.e., the *k^>^-prefixes*), and recognize the vertices at the ends of unipaths;

4) building *k^<^-prefix* set with the vertices at the ends of unipaths, and sorting all the *k^<^-prefixes* in lexicographical order;

5) merging the sorted tables of *k^>^-prefixes* and *k^<^-prefixes* to construct the *B_C_* representation of D.

All the steps are designed to guarantee constant RAM space cost and be affordable to the employed computers. Moreover, critical steps like (*k*+2)-mers sorting are also designed to accelerate the overall speed. A detailed illustration of transformation from (*k*+2)-mer set to unitigs (including GFA output) is in Supplementary Fig. 1

### 2.3 The enumeration of *k*-mers

In deGSM approach, (*k*+2)-mers of the input genome sequence(s) are enumerated at first. There have been many fast and memory-scalable *k*-mer counting tools which are suitable for this task, such as Jellyfish2 (Marçais and Kingsford, 2011), KMC3 (Kokot et al. 2017), etc. Jellyfish 2 is employed in the current version of deGSM. It is also worthnoting that Jelly-fish 2 is asked to run with limited RAM space.

The *k*- and (*k*+1)-mers at two ends of the input sequence(s) are also enumerated with deGSM itself. These *k*- and (*k*+1)-mers are sorted to remove redundant ones in later steps, and they are merged with the (*k*+2)-mers for further processing.

### 2.4 The sorting of *k*-mers

DeGSM implements an efficient parallel block sorting and multi-way merging to sort the enumerated (*k*+2)-mers. The whole dataset is divided into blocks to fit the size of the user-defined RAM space. Each block is subsequently loaded into memory and processed. For each block, deGSM sorts the k-mers in the three steps as following.

i) deGSM Splits the whole k-mer set into multiple subsets of the same size and assign them to each thread. In each thread, simply taking one *k*-mer’s 8-mer prefix as the hash address for rapid locating its bucket. Meanwhile, counting the numbers of k-mers in the buckets.

ii) deGSM allocates RAM space for each bucket according to its k-mer quantity in parallel. For one bucket, deGSM collects all of its k-mers from multiple threads to make k-mers in different buckets in lexicographic order.

iii) deGSM assigns the buckets to multiple CPU threads. Each thread performs a quick sorting on the *k*-mers of the corresponding bucket and outputs sorted *k*-mers to a temporary file.

When all the (*k*+2)-mer blocks are sorted, a multiple-way merging is implemented to make a sorted table for all the (*k*+2)-mers.

It is also worth noting that, for each of the (*k*+2)-mer, deGSM moves its first character to its end before sorting. This is useful for the next step to recognize the vertices at the ends of the unipaths. This can place (*k*+2)-mers and the same middle *k*-mers together after sorting. And the in- and out-edges of the vertices of the de Bruijn graph (i.e., *k*-mers) are easy to be investigated.

The *k*- and (*k*+1)-mers at two ends of the input sequence(s) are also sorted in a similar approach. During the sorting, each of the *k*- and (*k*+1)-mers is seen as pseudo (*k*+2)-mers, i.e., (*k*+2)-mers with an auxiliary character flanking at the two (for *k*-mers) or one (for (*k*+2)-mers) sides of the *k*- and (*k*+1)-mers, respectively. This indicates the corresponding vertex connects to an auxiliary vertex of the graph.

After sorting, deGSM removes all the redundant *k*- and (*k*+1)-mers and merges the sorted (*k*+2)-mers and pseudo (*k*+2)-mers, which are derived by the *k*- and (*k*+1)-mers, to construct a single sorted table (termed as *T_k+2_* in the later sections).

### 2.5 The recognition of the types of vertices

In *T*_*k*+2_, each line indicates a specific combination of a vertex (k-mer) and its in- and out-edges. All the vertices are then categorized in four types, i.e., ‘I’, ‘Y^+^’, ‘Y^-^’, and ‘X’, indicating the single in and single out, single in and multiple out, multiple out and single in, multiple in and multiple out vertices, respectively.

Due to the lexicographical order of the (*k*+2)-mers, all combinations of the same vertices with various edges are placed together in*T*_*k*+2_. So it is easy to recognize the types of the vertices by directly checking the combinations with the same first *k*-mers. DeGSM directly traverses *T*_*K*+2_ to mark the lines of *T*_*K*+2_ by *k*-mer types.

However, there are two exceptional situations as following (also refer to Supplementary Fig. 2 for a schematic illustration).

**Fig. 2:**
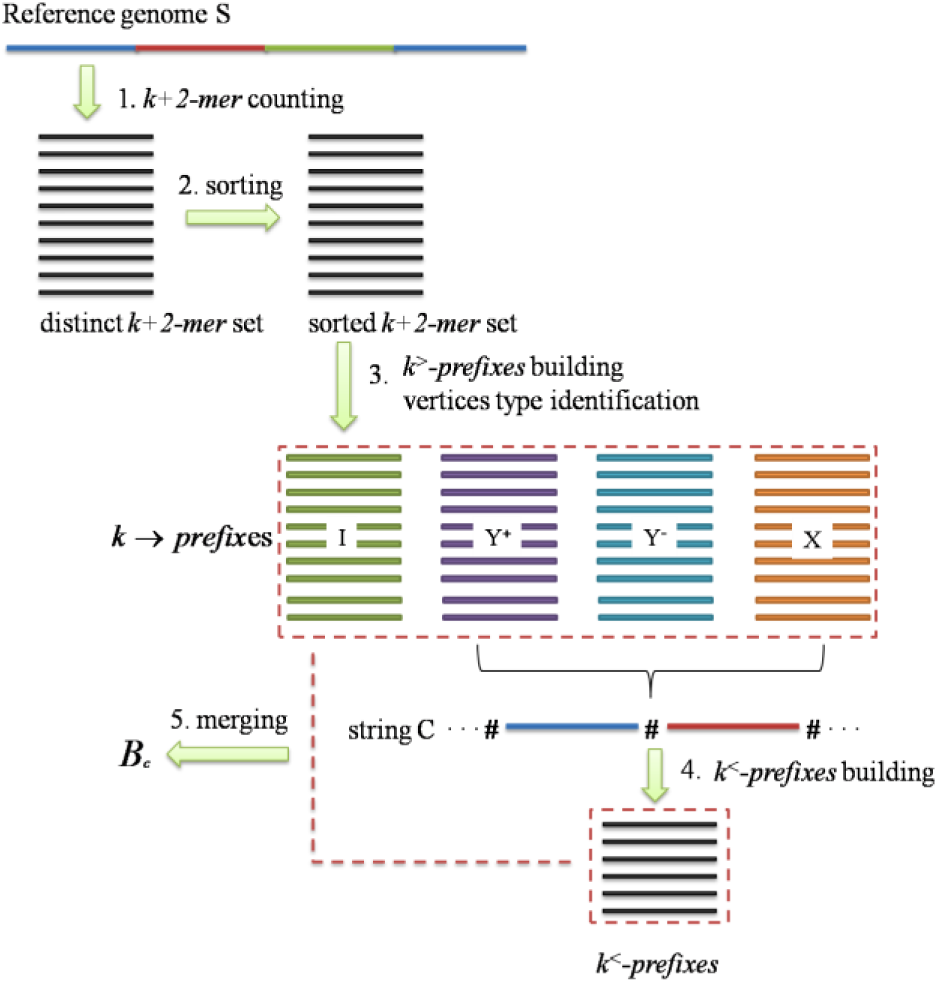
Flowchart of de Bruijn graph construction. Herein, various colored strips indicate repetitive fragments in S. The strips of different colors denote unipaths in C. The sorted k-mer sets are recorded in the external storage. The files that would never be used are to be deleted after each step.

1) For the multiple successor vertices of a Y^+^ type vertex, each of them is a starting vertex of a specific unipath, although it could be single in and single out. In this situation, deGSM marks such successor vertices as Y^-^ type, for the simplicity of later steps. Precisely, deGSM checks all the vertices having the same precursor. For each line of *T*_*K*+2_, the first character can also be seen as the edge linking to a precursor, and the substring from 2nd to *k*+1-th characters can be seen as the vertex itself. From this point of view, such successor vertices are also placed as neighbors in *T*_*K*+2_. In practice, deGSM divides *T*_*K*+2_ into blocks. The lines in each block have the same 2nd to *k*-th characters, indicating multiple vertices having the same *k*-1 prefix (Supplementary Fig. 2a). If some of the vertices have the same precursor, these vertices are just the successors which start unipaths. And deGSM marks the corresponding lines as Y^-^ type.

2) The multiple precursor vertices of a Y^-^ type vertex are also end vertices of various unipaths, and deGSM marks such vertices as Y^+^ type. DeGSM investigates the out-edges to find out these end vertices. As the lines corresponding to the precursor vertices have various first characters, they are not placed together in *T*_*K*+2_. However, their substrings from 2nd to *k*-th characters are same to each other and their lexicographical orders still remain (Supplementary Fig. 2b). Under such circumstance, deGSM partitions *T*_*K*+2_ into four blocks, and each of the blocks corresponds to the lines starting with a specific character (A/C/G/T). A 4-way traversing is implemented to find out the lines with same 2 to k substrings and the corresponding lines are grouped together to check the successors of the vertices (Supplementary Fig. 2c). If the investigated vertices have the same successor, they are determined as the precursors of a Y^-^ type vertex, and the corresponding lines are marked as Y^+^ type.

After type identification, deGSM merges the corresponding lines for each of the vertices to generate a sorted *k*-mer table. This straightforwardly forms the sorted table of *k^>^-prefixes*. Moreover, four sorted *k*-mer tables are virtually constructed, each corresponds to a specific *k*-mer type.

### 2.6 The construction of *k^<^-prefixes*

The sorted tables of Y^+^, Y^-^, and X type k-mers are used for formulating *k^<^-prefixes*. Since any permutation of the vertices at the two ends of the unitigs can be used to formulate a valid set of *k^<^-prefixes*, deGSM constructs a straightforward permutation. Precisely, assuming the sorted tables of Y^+^, Y^-^, and X type k-mers are respectively, 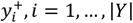, 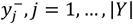, and where 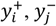 and *x_k_* are respectively the Y^+^, Y^-^, and X type k-mers, *i, j* and *k* respectively indicate the lexicographical orders of the three tables, and |*Y*| and |*X*| are respectively the numbers of Y^+^ and Y^-^ type k-mers, and the number of X type k-mers. DeGSM builds a simple permutation as following:

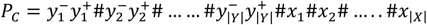

This can be also seen as a subsequence of C, and all the *k^<^-prefixes* consists of four categories of substrings of *P*_C_:

1) 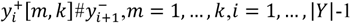, where 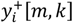 is the substring of 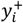 from the *m*-th character to the end of the *k*-mer;

2) 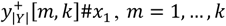 where 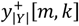 is the substring of 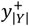 from the *m*-th character to the end of the *k*-mer;

3) 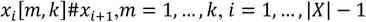 where *x_i_*[*m,k*] is the substring of *x*_i_ from the *m*-th character to the end of the *k*-mer;

4) 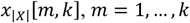, where *x_i_*[*m,k*] is the substring of *x*_|*x*|_ from the *m*-th character to the end of the *k*-mer.

A schematic illustration of the formulation of *k^<^-prefixes* is in Fig. 3.

**Fig. 3:**
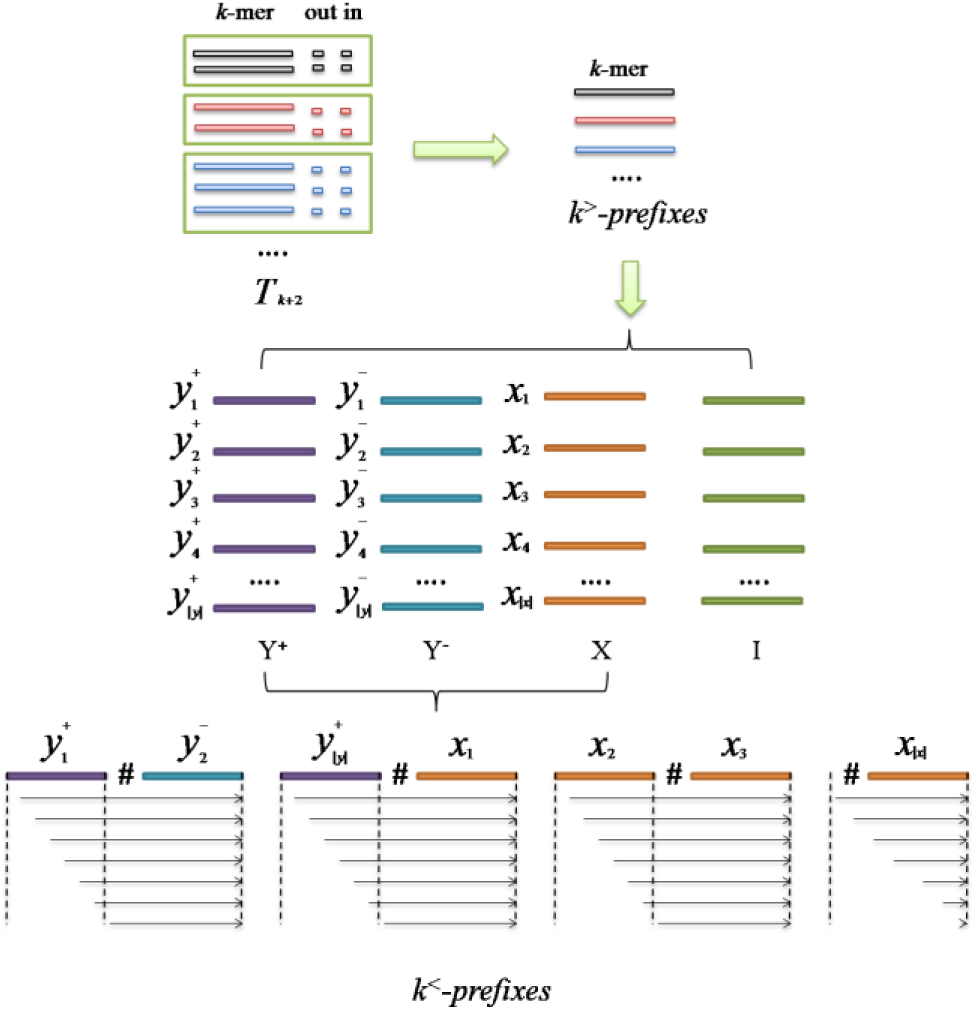
A schematic illustration of the formulation of *k^<^-prefixes*. The strips in various colors denote different *k*-mers. One line in *T*_*K*+2_ represents a certain k-mer and one of the combination of its in- and outedges. Sorted *k^>^-prefixes* can be derived by traversing *T*_*K*+2_. The *k^>^-prefixes* are categorized into four types (‘I’, ‘Y^+^’, ‘Y^-^’, and ‘X’), and *k^<^-prefixes* can be generated from the sets of ‘Y^+^’, ‘Y^-^’, and ‘X’ type *k-*mers.

All the *k^<^-prefixes* are constructed on-the-fly by simply traversing the whole sorted *k*-mer table, and they are recorded in a temporary file. After collection, deGSM constructs a sorted table of *k^<^-prefixes* by an approach similar to (k+2)-mer sorting.

### 2.7 The generation of the BWT of unitigs and the reconstruction of unitigs

DeGSM merges the sorted tables of *k^>^-prefixes* and *k^<^-prefixes* to determine the lexicographical order of all the suffixes of C string. As the (*k*+2)-mers are used in previous steps, the characters before the lexicographical order-determined suffixes are kept and the permutation of these characters are exactly *B_C_*.

Although unitig-BWT supports the query of any vertice and edge, the original unitigs are preferred in some applications. With *B_C_*, it is straight-forward to reconstruct all the unitigs with the LF-mapping. Each unitig sequence can be derived by backtracking from the last character ‘#’ on the bwt index, until the next ‘#’ is met. DeGSM provides a specific command for the conversion from unitig-BWT to original unitigs.

### 2.8 GFA format output

DeGSM provides the function to output in GFA (Graphical Fragment Assembly) format, to fulfil the requirements of the emerging graph-based sequence analysis tools, such as vg (Garrison et al., 2017). Therefore, each unitig is considered as a segment (S) in GFA. And the link (L) is represented by edge (overlapping k-1-mer) between two unitigs, which are stored during bwt backtracking. DeGSM retrieves unitgs to identify these connections by constructing outgoing k-mers of each unitig in bwt index (Supplementary Fig. 1).

### 2.9 *K*-mers filtering with specific abundance cutoff

*K*-mer filtration according to specific abundance cutoff is frequently used in sequence analysis tasks, such as genome assembly. For the filtered-out vertex, the out-edges of its prefix vertex or in-edges of its suffix vertex need to be modified, e.g., an end vertex can be changed to the internal vertex after its suffix vertex filtration out (Supplementary Fig.3a).

During vertex type recognition, the adjacent vertices in graph are often distributed into different data blocks, each of which cannot be loaded into memory simultaneously. DeGSM generates a novel *k*-mer set (called ‘pseudo *k-mer*’) as a signal to locate the prefix vertices or suffix vertices of the filter-out vertex (Supplementary Fig.3b). Meanwhile, deGSM sorts all pseudo *k*-mers and merge them with the reserved sorted *k*-mer set. After that, deGSM modifies vertex’s type identification from the gathered identical k-mers (Supplementary Fig.3c).

## 3 Results

DeGSM was implemented on a series of assembled genome sequence datasets and HTS datasets to assess its ability. Several state-of-the-art de Bruijn graph construction methods were also employed for comparison. All the benchmarks were conducted on a server with 2 Intel E5-2630v3 CPUs at 2.4 GHz (12 cores in total), 128 GB RAM and 48TB hard disk space (7200rpm RAID SAS hard disk drive with XFS File System, no SSD is used). In the benchmarking, 8 CPU threads were used as default, and deGSM was asked to run with upto 32GB RAM (most of the datasets are much larger than that) to assess its scalability (no such limit for other methods). For all benchmarked methods, the runtime of *k*-mer enumeration is excluded. All the command lines are available in Supplementary Notes.

### 3.1 Benchmarking on genome sequence datasets

DeGSM was assessed by two datasets from GenBank at first. Precisely, we downloaded the assembled contigs and scaffolds recorded in GenBank database (ftp://ftp.ncbi.nlm.nih.gov/genomes/genbank). All the domains (bacteria, viral, archaea, fungi, protozoa, invertebrate, plant, vertebrate_mammalian and vertebrate_other) are included, and the sizes of the two datasets are 305 Gbp (contigs) and 1.1 Tbp (scaffolds), respectively. The lengths of these two sets of sequences are orders larger than that of a single genome have been assembled, and the construction of de Bruijn graph on such large scale datasets is beneficial to large population genome or pan-genome analysis, as well as the assembly of very large genomes or metagenomes.

The results on the two GenBank datasets are shown in Table 1. DeGSM was asked to construct de Bruijn graph in various *k*-mer settings (*k* = 22, 30, 62 and 126, respectively), which are related to the requirements of various kinds of sequence analysis tasks. The total time costs, the time costs of various steps, the numbers of the vertices and the unitigs of the constructed graph, and the sizes of the temporary files are assessed. The results indicate that each of the constructed graphs consists of hundreds of billions of vertices, which are consistent with the lengths of the corresponding original sequences. The sizes of these graphs are one or two orders larger than the ones reported in previous studies (Birol et al., 2013; Chikhi et al., 2016). However, the tasks can be done with moderate memory footprint. Moreover, the time cost of the graph construction is also affordable, i.e., the graph construction for GenBank contigs can be done in about one day, and several days for GenBank scaffolds. Considering the graph size, memory footprint and time cost, deGSM is scalable and cost-effective.

**Table 1:**
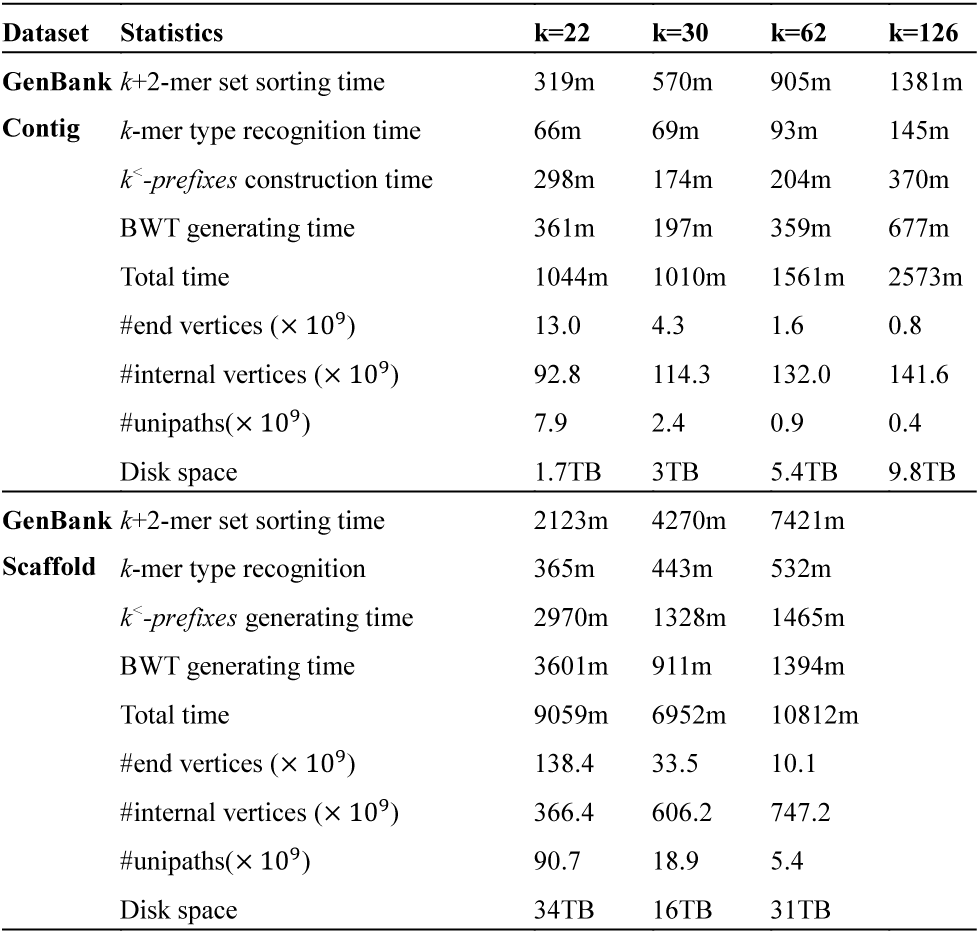
Statistics on GenBank Contig and GenBank Scaffold with various *k*-mer settings

DeGSM usually requires large hard disk space, as it produces several large temporary files to store the intermediate results of *k*-mer sorting and type recognition. For the two GenBank datasets, the temporary files occupied many terabytes hard disk space, moreover, the result on the GenBank scaffold dataset with k=126 is not shown due to that the temporary file size exceeds the hard disk space of the machine (48 TB). This could be a drawback of this approach, although a computer with large hard disk space is not hard to be available. Moreover, these files can raise intensive I/O operations, which affects the construction speed. It is observed that deGSM is faster with larger memory footprint, since more data can be loaded at once and I/O operations are reduced.

DeGSM was also implemented on the GenBank contig dataset with various multiple thread configurations (2, 4, 8, 12 and 16 threads). The time costs are in Table 2. The result indicates that time cost can be nearly 50% reduced with more threads. However, speedup attenuated due to that a few steps of deGSM are still hard to be operated in parallel way, such as the construction of *k^<^-prefix* and BWT generation. These steps have intensive multiple-way merging and I/O operations and are still non-trivial to make good parallel implementations.

**Table 2:**
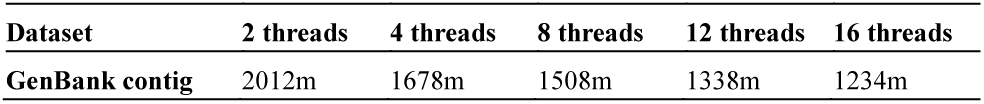
Runtimes with various numbers of threads for the dataset of GenBank Contigs (k=55)

We tried to implement three other state-of-the-art de Bruijn graph construction approaches, TwoPaCo, MEGAHIT and a BWT-based method (Baier et al, 2016), on the two GenBank datasets for comparison. However, all the three methods collapsed, mainly due to the fact that the RAM usage exceeds the 128 GB RAM space of the machine. To make a fair comparison, we benchmarked the methods on a smaller dataset (termed as GenBank-small) that all the methods can be successfully run. This dataset built by randomly selecting 61 files of the GenBank contigs, and its size is 3.1 Gbp similar to a human genome.

The result of GenBank-small dataset is shown in Table 3. Only the result of k=31 is shown, as TwoPaCo collapsed with larger k parameter setting again, due to its large memory footprint. It is observed that deGSM has outstanding speed, that is several times faster than TwoPaCo and the BWT-based method and outperforms MEGAHIT.

**Table 3:** Runtimes of the small GenBank contig dataset (k=31)

| Dataset | Statistics | deGSM | TwoPaCo | MEGAHIT | Bwt-based method |
| --- | --- | --- | --- | --- | --- |
| GenBank-small | Memory | 16GB | 74GB | 79GB | 8G |
|  | Time | 18m + 3m | 120m | 34m | 104m |

Considering the ability to construct very large graphs, small memory footprints and relatively fast speed, deGSM is scalable and useful to handle very large genome sequence datasets.

### 3.2 Benchmarking on HTS datasets

To assess the ability of deGSM to construct de Bruijn graph for HTS data, we implemented deGSM on a high coverage HTS dataset of the 20 Gbp *Picea abies* genome. The dataset (SRA accession: ERP007725) consists of 94.9× 109reads in 52-202 bps produced by Illumina platforms. Two *k*-mer settings were used (k = 29 and 53) in the assessment to mimic commonly used configurations in de novo assembly.

*K*-mers are often filtered in genome assembly, since *k*-mers having low quality bases or low abundance are usually false positive ones produced by sequencing errors. The rules of the filtration are of the tradeoffs between sensitivity and specificity. And various rules have been used in proposed de novo assembly approaches. In this assessment, two rules were respectively used for the two *k*-mer settings, i.e., i) for k=29, a k-mer is filtered out if it occurs less than 3 times; and ii) for k=53, a k-mer is filtered out only if there is at least one base having very low Phred Quality Score (Q<3), which indicates a failure in sequencing. These two rules are quite conservative, i.e., they focus on keeping more true positive k-mers, but prevent an explosive number of k-mers which may make the temporary files be out of hard disk space. The rules are similar to that of some existing de novo assembly approaches, which are designed to achieve high sensitivity in initial steps.

The results on the *Picea abies* dataset are shown in Table 4. The sizes of the constructed graphs with *k*=29 and 53 are on the same order of the GenBank contig and scaffold datasets. It is also worth noting that, the number of the vertices of the two graphs (69.9 ×10 9 and 402.8 ×10 9, respectively) are much higher than the length of *Picea abies* genome (about 20 Gbp). Moreover, the number of 53-mers is about 6 times larger than that of 29-mers. This indicates that there exist serious sequencing errors and many false positive *k*-mers still remain after the filtration, which makes explosive growth on graph size. This could be difficult for many current de novo assembly approaches as the memory footprint may drastically increase. Meanwhile, such case can be handled by deGSM well.

**Table 4:**
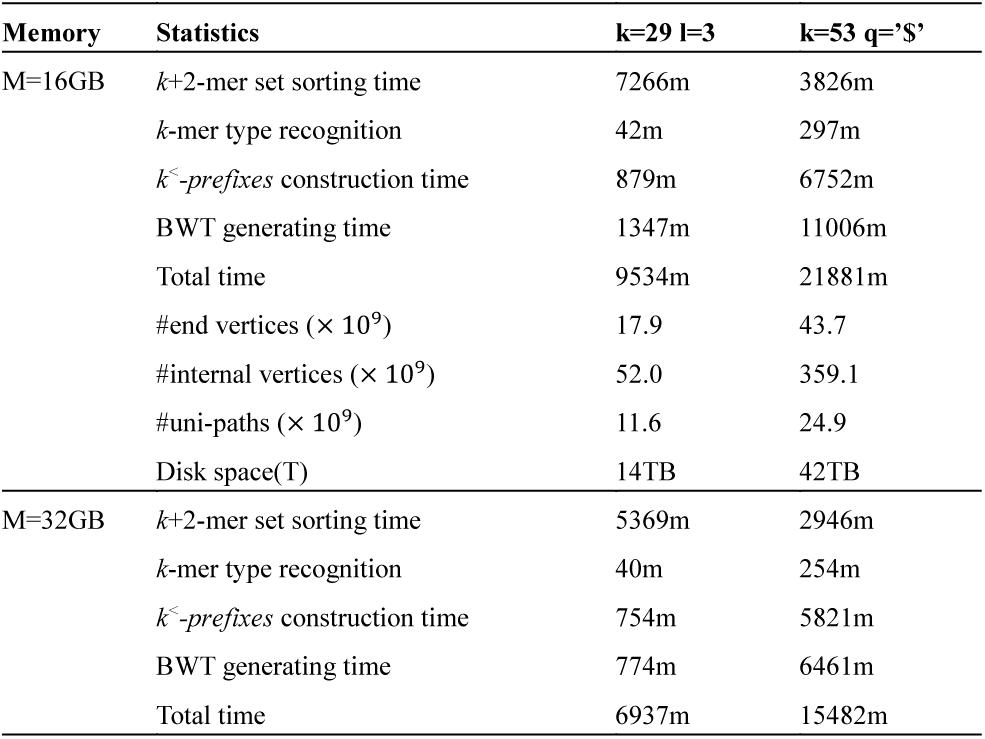
Statistics on Picea abies with respect to different memory foot-prints

The time costs of HTS datasets are higher than those of GenBank datasets. This is mainly caused by the large amount of false positive *k*-mers. Most of the false positive *k*-mers can produce extra branches, which can make numerous end vertices and very short unitigs. These end vertices greatly increase the time costs of some steps of deGSM, since the merging of the *k*-mers raises intensive I/O operations. However, this performance degradation can be mitigated with less false positive k-mers, since there would be less extraordinary short unipaths and end vertices and the time cost to handle these vertices can be greatly reduced. This is partially suggested by another benchmarking on a simulated *Picea abies* HTS dataset with lower sequencing error (shown below), that the time cost largely decreases with less false positive *k*-mers.

In practice, the number of false positive *k*-mers strongly depends on the quality of HTS data and the adopted filtration rule, i.e., better sequencing quality can greatly reduce false positive *k*-mers, meanwhile, many advanced filtration methods have been proposed, which can also effectively filter such *k*-mers out. With less false positives, deGSM can still efficiently construct de Bruijn graph, even if the number of true positive *k*-mers is huge (considering that of the two GenBank datasets). Under such circumstance, deGSM can be also beneficial to HTS data analysis tasks such as de novo assembly, especially with effective *k*-mer filtration approaches.

DeGSM is compared with two state-of-the-art de Bruijn graph construction methods for HTS datasets, BCALM2 and MEGAHIT, on two simulated HTS datasets. These two methods are out of RAM space for the real *Picea abies* HTS dataset. In precise, we used ART (Huang et al., 2012) to simulate two HTS datasets. One (termed as PA-sim) is a 70X *Picea abies* HTS dataset with moderate sequencing error rate. And the other one (termed as GenBank-sim) is a 6X simulated HTS dataset from a part of GenBank contigs (10.8GB in total) with low sequencing error. The results on these two datasets are listed in Table 5.

**Table 5:** Runtimes of the simulated datasets from Picea abies genome and GenBank contigs (k=51)

| Dataset | Statistics | deGSM | BCALM2 | MEGAHIT |
| --- | --- | --- | --- | --- |
| PA-sim | Memory | 32GB | 79GB |  |
|  | Time | 3660m + 67m | 1158m |  |
| GenBank-sim | Memory | 16GB | 41GB | 95GB |
|  | Time | 450m + 15m | 270m | 294m |

On PA-sim, *k* is set to 51 as this *k*-mer size is similar to those commonly used settings in de novo assembly. Meanwhile, only the *k*-mers occurred less than 3 times were filtered out. DeGSM is slower but still affordable and comparable to that of BCALM2. False positive *k*-mers are still the main issue causes the slowdown of deGSM, i.e., many false positive *k*-mers still remain after filtration, although the sequencing error is in moderate level. However, this is not as serious as that of the real *Picea abies* HTS dataset, i.e., the proportion of false positives is lower, and the graph can be constructed with obviously faster speed. MEGAHIT collapsed on PA-sim due to out of memory space.

To further investigate the impact of false positives on the graph construction, deGSM is implemented on GenBank-sim, which has a low sequencing error. Both of BCALM2 and MEGAHIT can construct the graph with the 128 GB RAM space. The time cost of deGSM is much closer to that of BCALM2 and MEGAHIT. This indicates that the speed of deGSM can be substantially improved with lower sequencing error, due to less false positives.

The results of BCALM2 and MEGAHIT indicate the advantage of inmemory approaches. The I/O operations can be greatly reduced. It is also easier to implement parallel operations, such as the thread-safe queues and Minimal Perfect Hash Function (MPHF) (Cormen, 2009) of BCALM2, and the in-memory parallel sorting (CX1 algorithm) (Liu et al., 2014) of MEGAHIT, to accelerate the speed. However, the data structures used by in-memory approaches may require a prohibitively large RAM space, which is a bottleneck to handle large datasets.

## 4 Discussion

Large scale sequence analysis is promising in many cutting edge genomic studies nowadays. With the explosive growth of HTS data and assembled genomes, there is a high demand to analyze sequences in Tera bp scale. As a fundamental data structure, de Bruijn graph may play an important role. However, it is still lack of highly scalable de Bruijn graph construction approaches which can well handle such large sequences, especially, the RAM space of computer is usually being limited in practice. As it is hard to unlimitedly increase RAM space, the lack of effective de Bruijn graph construction approaches could make it a bottleneck to many forthcoming sequence analysis tasks, especially the size of the sequence to be handled is ever-increasing.

Herein, we propose a highly scalable de Bruijn graph construction approach, deGSM, which can well handle very large sequences. Like other suffix-trie-based de Bruijn graph construction approach, deGSM takes advantage of the relationship between suffix trie and de Bruijn graph. Taking advantage of the novel organization and efficient external sorting of *k*-mers, deGSM can effectively build the BWT of the unitigs, i.e., the unitig-BWT representation of de Bruijn graph. In theory, deGSM is capable of constructing any size de Bruijn graph with any given RAM space. In practice, the implementation of deGSM fully considers the configurations of commonly used computers, to achieve a balance between speed and RAM usage. The benchmarking results on a series of very large genome sequences and HTS datasets demonstrate that deGSM can construct very large de Bruijn graph (one or more orders larger than that of previous studies) in affordable time, with only a moderate hardware configuration. This could be very beneficial to break through the bottleneck of large de Bruijn graph construction.

A drawback of deGSM is that the time costs of the multiple way merging-related steps are quite large, especially when there are a large proportion of false positive k-mers caused by serious sequencing errors in HTS data. Due to the serial processing and intensive I/O operations, it is non-trivial to reduce the time greatly. However, this can be mitigated with less false positive k-mers. In this situation, an advanced k-mer filtration method could be very helpful, since it can reduce false positive k-mers while keeping true positive ones. Meanwhile, ubiquitously used SSD could be also helpful as it can greatly improve the speed of I/O operations on external storage.

It is also worth noting that, a machine with large RAM could be still required to transform the BWT string into untigs, as the entire BWT string need to be loaded into RAM. However, this requirement is not hard to fulfill, since the BWT string is a very compact de Bruijn graph representation and its size is not very huge, e.g., 87 Gigabytes for the GenBank contig dataset. This can be handled by a machine with large RAM space, while time cost of the transformation is linear to the length of the BWT string which is not large. Moreover, it only needs to execute once to obtain all the original unitigs. And the unitigs could be further re-used in various ways with less RAM space. For example, the unitigs could be compressed with advanced compression approaches to make a more compact representation of the de Bruijn graph. And for some tasks, like scaffolding, pangenome analysis, metagenomics HTS read classification, etc., it is also possible to load only a portion of unitigs instead of the entire graph into memory.

Overall, with its scalability, deGSM is a promising tool for the de Bruijn graph construction of large sequences. It may have enormous potentials in large scale genomic studies.

